# Estimating the timing of HIV infection from unmutated sequences

**DOI:** 10.1101/2020.11.28.402271

**Authors:** Alec Pankow, Murray Christian, Natalie Smith, Daniel Sheward, Ben Murrell

## Abstract

For HIV, the time since infection can be estimated from sequence data for acutely infected samples. One popular approach relies on the star-like nature of phylogenies generated under exponential population growth, and the resulting Poisson distribution of mutations away from the founding variant. However, real-world complications, such as APOBEC hypermutation and multiple-founder transmission, present a challenge to this approach, requiring data curation to remove these signals before reasonable timing estimates may be obtained.

Here we suggest a simple alternative approach that derives the timing estimate not from the entire mutational spectrum but from the proportion of sequences that have no mutations. This can be approximated quickly and is robust to phenomena such as multiple founder transmission and APOBEC hypermutation. Our approach is Bayesian, and we adopt a conjugate prior to obtain closed form posterior distributions at negligible computational expense.

Using real data and simulations, we show that this approach provides accurate timing estimates and credible intervals without the inconvenience of data curation and is robust to complicating phenomena that can mislead existing approaches or cause them to fail entirely. For immediate use we provide an implementation via Google Sheets, which offers bulk analysis of multiple datasets, as well as more detailed individual-donor analyses. For inclusion in data processing pipelines we provide implementations in three languages: Julia, R, and Python.

## Introduction

HIV prevention efficacy trials can leverage accurate inference of time since infection (hereafter referred to as “infection time”) in order to identify correlates of protection. ^1^ While clinical diagnostic staging can inform timing estimates, there is particular interest in using viral sequence data from early in infection for this purpose. For HIV, roughly 80% of infections are homogeneous, initiated by one distinct founding strain ^2–4^, suggesting a strong transmission bottleneck. While the exact mechanism of the transmission bottleneck is unknown, there is evidence that route of transmission is associated with increased odds of observing multiple founder infections. ^3,5,6^ These estimates were based on relatively shallow sequencing, and it is possible that the deployment of higher throughput sequencing will reveal that the transmission of multiple founder variants occurs even more frequently.

Following the establishment of the initiating founder strain(s), HIV typically grows rapidly and exponentially ^7^. This leaves an imprint on the resulting phylogeny: the tree is largely “star-like” with all lineages coalescing near the most recent common ancestor (MRCA). ^8^ It has been observed that acute HIV infections with a homogeneous strain typically follow this pattern. ^2^ The diversity of this acute-infection virus population has been found to increase roughly linearly with time, motivating the development of several methods for diversity-based infection time estimation. ^9–14^ One popular method is Poisson-Fitter, ^15^ a maximum likelihood approach which estimates infection time from the distribution of pairwise Hamming distances between aligned sequences, under the assumption that the number of mutations from the founder to each observed sequence follows a Poisson distribution. Violations of this assumption have been attributed to APOBEC mediated hypermutation, the transmission of multiple variants, the onset of immune selection, or stochastic early mutations. ^15,16^

Hypermuation via host APOBECs (apolipoprotein B mRNA-editing catalytic polypeptides) introduces G to A mutations into the HIV genome by cytidine deamination of the negative strand cDNA. ^17–19^ While typically understood as an “all-or-nothing” phenomenon associated with defective viral Vif, ^20,21^ several studies have demonstrated that more subtle, sub-lethal levels of can occur in *in vitro* experiments. ^22,23^ Additionally, studies using Poisson-Fitter timing estimates have noted that removal of APOBEC-targeted sites restores a Poisson distributed mutational spectrum. ^2,24^

For multiple founder infections, prior to the onset of immune selection (Fiebig stage I and II) and recombination, a theoretical population resembles a collection of star-like phylogenies - one for each founder - each with the same root-to-tip distances. If the founder strains can be discriminated, and all sequences assigned to the correct founder, then each founder can be modelled separately to obtain independent estimates of *λ*. In practice, the difficulty of such splitting can range from trivial, to difficult, to nearly impossible. Multiple variant transmission from acute donors is known to occur, and may be more prevalent than previously appreciated. ^25^ It is particularly unclear in this scenario how to distinguish closely-related founder variants that diverged in the donor population from stochastic early mutations separating variants that diverged in the recipient population. A variety of founder identification methods of varying complexity exist, ^26–28^ and do not uniformly agree.

Both APOBEC and multivariant transmission are examples of processes that inflate the average pairwise distance between sequences and cause violations of the Poisson assumptions that Poisson-Fitter relies upon. These can be remedied by curation, removing APOBEC hypermutated sequences or sites and grouping sequences by founder before attempting to estimate the time since infection. These curation steps are non-trivial however, and both introduce a substantial effort burden on the user and potentially multiple user-level decisions for each infection time estimate, which may be statistically problematic, depending on how these estimates are used. ^29^

Here we describe ZFitter (for “Zero-Fitter”) which, like Poisson-Fitter, aims to estimate infection time from HIV sequence datasets, but is designed to be more robust to the Poisson violations that are routinely observed in HIV sequence datasets.

## Methods

### Inference

ZFitter begins with the observation that, for most datasets sampled from acute infection, there are multiple sequences that are identical to each other, typically representing variants that have not mutated away from the founder. Like Poisson-Fitter, ZFitter assumes that mutations are Poisson distributed, with parameter *λ*, which we can subsequently relate to “time” through a known mutation rate. Under this Poisson assumption, the number of sequences with no mutations, *s*, among a fixed total of *N* sampled sequences, is Binomial distributed with probability mass function

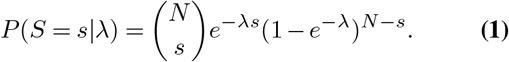

ZFitter is Bayesian in nature, and we introduce a two-parameter family of conjugate prior distributions for the above likelihood, denoted 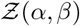. The density functions are defined for all *α, β* > 0, and all *λ* > 0, as

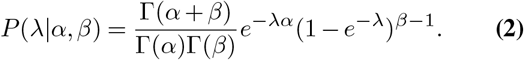

Assuming a 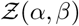 prior over *λ*, the posterior distribution is 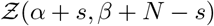. Figure 1A shows Poisson distributions (and the expected proportion of sequences with no mutations) for varying *λ*. And using our standard prior of *α* = 0.3*, β* = 1 (which we use throughout), figure 1B displays example posterior distributions corresponding to a range of *N* and *s* counts. We consider cases where *s* = 0 to be “inestimable” by ZFitter, where all we can do is provide a lower-bound on the infection time.

**Fig. 1.**
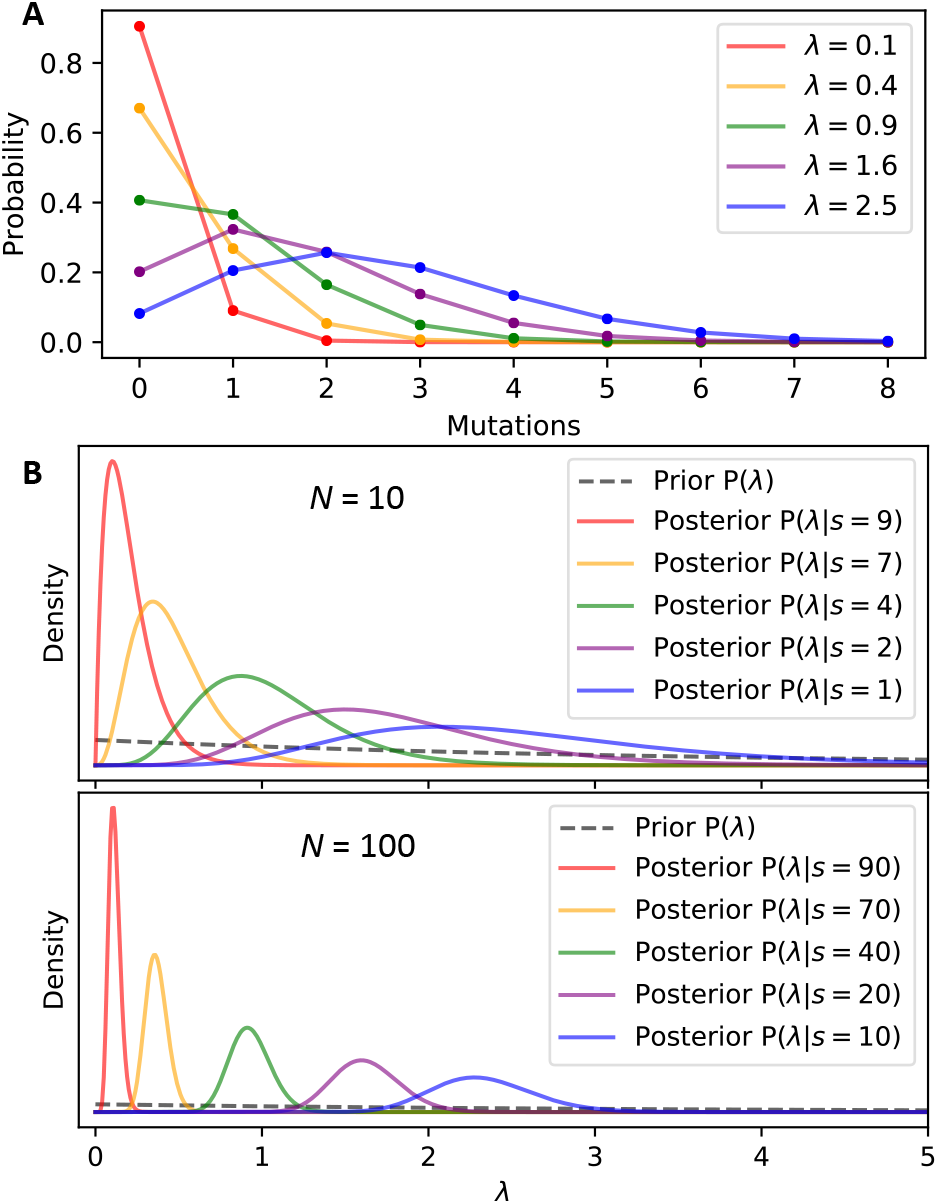
Relationship between mutation rate and the number of sequences with no mutations. **A**. Poisson distributions for varying *λ*. Our “ZFitter” approach uses only the proportion of sequences that have zero mutations. **B**. Posterior distributions over *λ* for two different sample sizes (N =10, and N =100), each with five different counts of unmutated sequences. These posterior distributions over *λ* can be translated to time through an *apriori* known or estimated mutation rate.

Our parameterisation of the family is motivated by the observation that 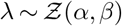 if and only if *e*^−*λ*^ *Beta*(*α, β*). Using this, the quantile function for a 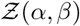-distributed random variable, *F*^−1^(*q*), can be expressed in terms of the standard quantile function for a *Beta*(*α, Beta*)-distributed variable, 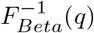, as

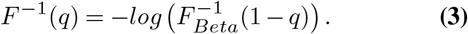

All 95% Bayesian “Credible Intervals” (CIs) presented here are *F*^−1^(0.025) to *F*^−1^(0.975). This approach has a number of computational and statistical consequences. Computationally, we can approximate *s* relatively well with the number of identical sequences, which is trivial to compute from sequence datasets. A multiple sequence alignment is not even required for this. Secondly, given chosen prior parameters and *s* and *N* counts, the conjugacy above provides closed-form expressions for posterior medians, and any required posterior credible intervals.

Statistically, ZFitter involves a trade-off. On the one hand, not all the information in the sequences is exploited, as the full mutational spectrum is ignored, attending to only the zero-valued mass. Under perfect Poisson assumptions, ZFitter should thus provide less precise and less confident estimates of time since infection. On the other hand, this renders ZFitter far more robust to particular assumption violations that frequently occur in real-world data. Key among these are APOBEC-mediated hypermutation, which dramatically inflates the mutation rate (real or apparent) of a small number of sequences, and multiple founder infections. Both can cause a dramatic shift in the distribution of pairwise distances, but will only minimally affect the proportion of completely unmutated sequences. We demonstrate this robustness through simulation.

### Implementation

The approach described here is so computationally trivial that it can be implemented in a simple spreadsheet. Indeed, we offer a public “Google Sheets” implementation: https://bit.ly/3pOsa2a. This offers two kinds of functionality: i) a bulk processing option, where counts of sequences (total, and non-singletons) are input for a large number of datasets, and infection estimates and CIs are provided for each, and ii) a single-dataset processing option, where either counts are entered, or sequences are pasted in directly, and both the estimates, and a plot of the prior and posterior distributions are displayed. For convenient incorporation into computational pipelines, simple implementations in Julia, Python and R are included in Supplemental S1-3.

### Simulations

To generate simulated sequences from a founder *env* sequence, an HKY85^30^ substitution rate matrix, *Q*, was used with a transition to transversion ratio of 4.5. Equilibrium frequencies were calculated from an HIV dataset derived from a single donor ^31^: *π*_*A*_ = 0.34, *π*_*C*_ = 0.17, *π*_*G*_ = 0.23, *π*_*T*_ = 0.24. The *Q* matrix was scaled to yield an average rate of 1.19e-5 substitutions per site per day; the daily internal rate used by Poisson-Fitter. ^28^ All individual simulations, unless otherwise noted, consisted of 100 sequence observations and time, *t*, was varied from 5 to 80 days. Mutations are sampled from *P* = *e*^*Qt*^.

### Star-like, single founder

Infections were modelled as star trees with all observed sequences equidistant from the root. Branch lengths were set equal to the time (in days) since infection.

### Star-like, dual founder

Two separate founder sequences were sampled from the “source” population, derived from one timepoint of a longitudinally-sampled donor from a primary infection cohort. ^31^ Founder frequencies were drawn from a Multinomial distribution, which itself was drawn from a Dirichlet distribution with a concentration parameter of 1. Sequences from each founder were then mutated as in the single-founder case.

### APOBEC-mediated hypermutation

APOBEC rates were simulated from a mixture of Gamma distributions:

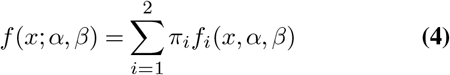

with prior weights *π*_1_ = 0.9, *π*_2_ = 0.1 and gamma parameters *α*_1_ = 1, *β*_1_ = 1*e* − 4, *α*_2_ = 10, and *β*_2_ = 0.006. This models a situation where 90% of the sequences are only weakly affected by APOBEC activity (on average only 1.2% of this 90% have any APOBEC mutations at all), but 10% of sequences have more substantial APOBEC effects, mimicking those observed in more severe APOBEC-affected datasets. Context-dependent APOBEC effects were introduced proportional to their empirical occurrence across trinucleotide contexts estimated from a control dataset. Fig ED1 shows the frequencies of APOBEC mutations per sequence introduced by this scheme.

## Results

### Simulated

Figure 2 summarises the performance of ZFitter and Poisson-Fitter on simulated sequence datasets. For single founder infections, where Poisson assumptions are clearly satisfied, estimates from both methods closely tracked the simulated infection time (Figure 2A). For single founder infections with APOBEC mediated hypermutation, ZFitter performed similarly to post-curation Poisson-Fitter with all APOBEC positions scrubbed from the alignment, and was significantly m ore a ccurate t han P oisson-Fitter without any curation (Figure 2B). In comparison to true infection time, both curated Poisson-Fitter and Zfitter estimates for the lowest range of infection times were biased slightly upward. For ZFitter, this is because a high percentage of APOBEC-mutated sequences presents a non-negligible influence on the number of completely unmutated sequences for smaller *t*, but matters relatively less when *t* is large. It is not yet clear why post-curation Poisson-Fitter exhibits this behavior as well. For both dual founder simulations, ZFitter was able to infer reasonable estimates of infection time from all combined sequences, closely matching curated Poisson-Fitter, which, here, relied on perfect knowledge of how to group the sequences by founder, and was estimated for the largest lineage only (Figure 2C,D).

**Fig. 2.**
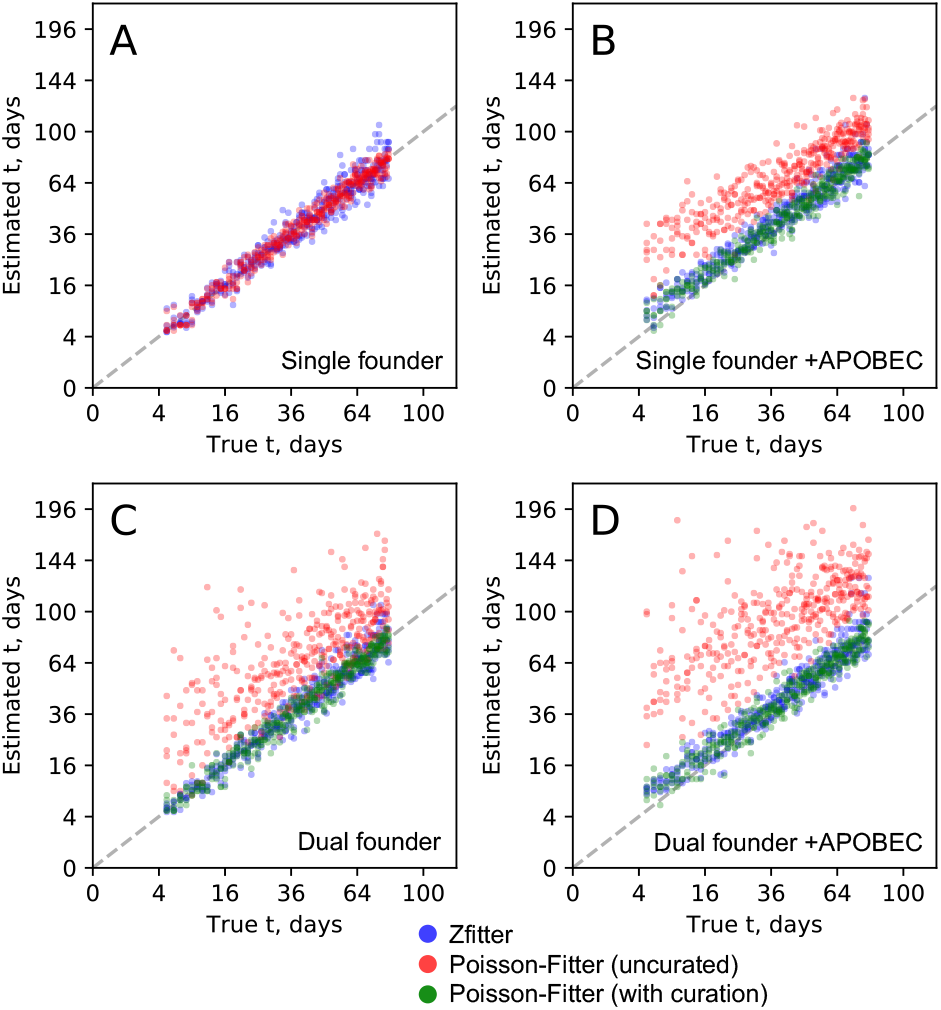
ZFitter and Poisson-Fitter estimates for simulated datasets. Simulated (“True”) infection times ranged from 5 to 80 days. A 1:1 line is included for reference. **A.** Star-like, single founder. Zfitter (blue) and Poisson-Fitter estimates (red) are shown, with no Poisson-Fitter correction. **B.** Star-like, single founder with simulated APOBEC hypermutation. Both Poisson Fitter estimates for all sequences (red) and with APOBEC positions removed from the alignment (green) are shown. **C.** Star-like, dual founder. Both Poisson Fitter estimates for all sequences (red) and the largest homogeneous virus lineage (green) are shown.**D.** Star-like, dual founder with simulated APOBEC hypermutation. Both Poisson Fitter estimates for all sequences (red) and the largest homogeneous virus lineage with APOBEC positions removed from the alignment (green) are shown.

### Empirical

ZFitter estimates were obtained from 130 Sanger SGA datasets from two published studies on acute HIV infection. ^2,6^ For an in-depth description of the results of each dataset and associated metadata, see Table S1. Figure 3 displays the correlation between ZFitter and Poisson-Fitter for the 109 out of 130 datasets with a published Poisson-Fitter estimate. As expected from the simulations, Poisson-Fitter and ZFitter largely agree when the distribution of pairwise Hamming distances suggest that the Poisson assumptions are not violated. Also as expected from the simulations, violations of the Poisson model tend to produce Poisson-Fitter estimates which are skewed toward longer infection times. This is particularly apparent in case of multiple founder infection (as adjudicated in the original publications). The median ZFitter estimated infection time for all multiple founder infections is 27 days, while the median Poisson-Fitter estimate is 136 days. For 4 datasets (Z03, SC42, Z29, and 4013291), all sequences were unique, producing elevated and diffuse ZFitter posterior distributions.

**Fig. 3.**
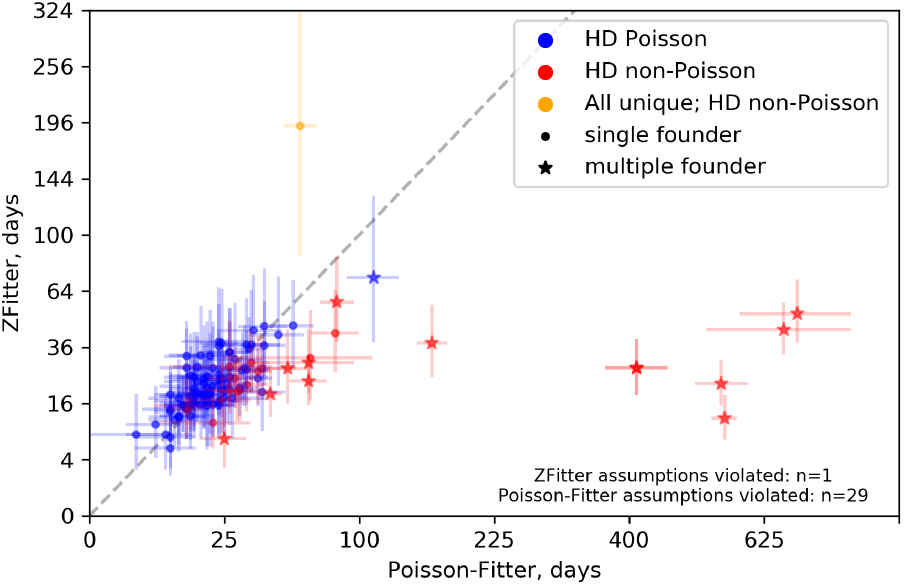
ZFitter and Poisson-Fitter estimates for 109 Sanger SGA datasets. Horizontal and vertical rules represent the 95% CI for both estimates. A 1:1 line is included for reference. Points are colored by whether or not the Poisson assumptions were supported by the distribution of pairwise Hamming Distances (HD - groups “HD Poisson” and “HD non-Poisson”) and whether or not all observed sequences were unique.

Figure 4 displays ZFitter estimates for all 130 datasets grouped by Fiebig stages. ^32^ Timing estimates from infections with Fiebig stages II and III differed significantly from infections in Fiebig stage V, but all other comparisons were not significant (Mann-Whitney U Test). The relatively fast clinical progression through stages II, III, and IV could explain part of why their median ZFitter estimates are so similar. Stages II, III and IV have typical duration times of five, three, and six days, respectively. ^32^ Stage V is longer, with a median duration of 70 days.

**Fig. 4.**
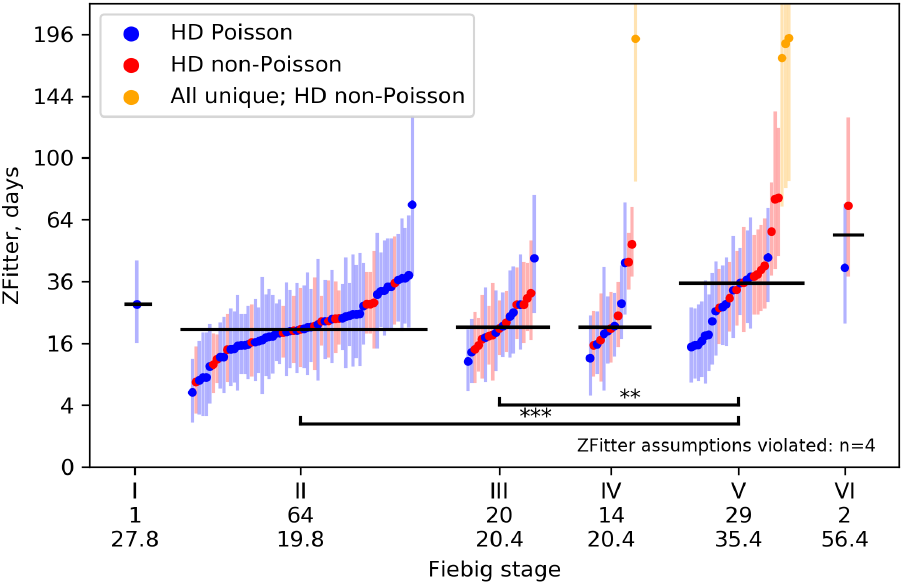
ZFitter estimates by Fiebig stage for 130 SGA datasets. Vertical rules represent the 95% CI for each estimate. Horizontal rules for each Fiebig stage indicate the group median. The number of datasets for each stage and the median estimate (in days) is included in the x axis label. Points are colored as in Figure 3.

## Discussion

Accurately estimating the date of HIV infection is a critical parameter for on-going clinical trials aimed at HIV prevention, where the knowledge of the titer or concentration of inhibitors at the time of infection is required to determine the correlates of protection. ^1^ In a recent study, Poisson-Fitter, a maximum likelihood approach which models the distribution of nucleotide mutations in a acute infection, yielded more accurate, more precise and unbiased estimates for the time of infection than did coalescent phylogenetic models implemented in BEAST. ^24,33^ However, violations of the Poisson-Fitter model assumptions of star-like phylogeny and a Poisson-distributed accumulation of mutations by various phenomena, including APOBEC-mediated hypermutation, immune mediated selection, recombination, or multiple-founder transmission can mislead Poisson-Fitter’s estimates, requiring manual data curation and iterative Poisson fitting. Here, we developed a method which is more robust to these complications by only considering if a sequence has mutated away from the founding strain. This approach does not leverage all the information in the sequences and will therefore be less precise when mutations are truly Poisson distributed. However, this is a potentially beneficial trade-off, as many assumption violations will dramatically skew the distribution of pairwise distances but not substantially alter the proportion of unmutated sequences. The performance of Zfitter on simulated datasets (Figure 2) demonstrates its comparable performance to Poisson-Fitter when infections are homogeneous, as well as ZFitter’s robustness to both APOBEC mediated hypermutation and multiple founder infection, which require additional curation for reasonable Poisson-Fitter estimates. The simulated level of APOBEC only induces small bias in estimated infection time for low *t*. This is in contrast to Poisson-Fitter, which is relatively sensitive to these sequences as they induce strong perturbations to the distribution of pairwise Hamming distances and skew *λ* upwards. And even when APOBEC sites are removed, Poisson-Fitter appears to have the same bias at low *t* as ZFitter.

To investigate ZFitter’s behavior on real data from acute HIV infection, we processed 130 Sanger SGA datasets from published studies with Poisson-Fitter estimates of infection time. ^2,6^ ZFitter and Poisson-Fitter estimates were largely consistent when there was a good Poisson model fit, albeit with larger uncertainty than observed in our simulations (Figure 3). This was at least partly due to lower sampling depth in available datasets (median=28). In a similar manner to the dual-founder simulations, in instances of multiple founder infection ZFitter produced timing estimates more consistent with acute infection when run on all available sequences.

There are several caveats to our approach which are important to discuss. As all of our signal comes from the number of sequences which cluster together, estimates made on any dataset where all sequences are unique or the ratio of *s* to *N* is very small should be treated with caution. One biological process that can fragment the founding virus lineage is early T cell mediated selection. This is predominantly positive/directional selection by CD8 + T cells narrowly focused on a small number of epitopes and is known to occur relatively early in infection associated with initial control of plasma viral load. ^34–37^ We have not investigated the effect of such a process on our inference since we do not know how to appropriately simulate it, but we note that for real, single-founder datasets, our estimates did not appear to be biased either way compared to Poisson-Fitter. Therefore, at the very least we are not especially affected by this process.

Another potential source of error is recombination between distinct founder strains. Recombinant sequences are frequently detected in acute infection when initiated by two or more founding strains. ^2,5,6^ When they contain unique break-points, these sequences inflate the number of u nique sequences in the data. We do not explicitly filter for recombinant sequences currently in ZFitter, as most standard methods for recombination detection either require specification of founding lineages in advance (RAPR) ^38^ or require an amount of phylogenetic signal which may be lacking in instances of multiple founder infection from a low diversity source (RDP4). ^39^ However, we note that the effect of recombination on *s* is similar to that of APOBEC: even extreme APOBEC hypermutation of a sequence will reduce *s* by one, just as would that sequence harbouring a single mutation, or a sequence being the recombined offspring of two other sequences. This is to say that, as long as the *per-sequence* probability of at least one mutation is substantially higher than the probability of recombination or hypermutation, ZFitter’s estimate should remain relatively robust to such processes.

ZFitter may open the door for new strategies for identifying founder variants and grouping sequences into founder clades. For low-diversity multivariant founder infections, the founder identification problem suffers from a chicken/egg issue. It can be difficult to split a dataset into founders without knowing the amount of post-infection divergence. But estimating the post-infection divergence, especially with methods such as Poisson-Fitter, require that datasets are already split into founding clades. By providing robust estimates of post-infection divergence without any curation, simply relating the number of unmutated sequences to the expected divergence, ZFitter’s *λ* estimates may be useful as inputs to founder clustering algorithms, which we will explore in future work.

Here we have shown that the performance of ZFitter on real and simulated data supports its further investigation as a timing estimator for sequences from acute HIV infection. The method is trivial to implement and is designed to require no sequence curation to obtain timing estimates. Where extensive sequence curation is feasible, ZFitter should provide a valuable supplementary method to existing approaches such as Poisson-Fitter, allowing a consistency check by comparing the uncurated ZFitter estimate to the curated Poisson-Fitter estimate. Where curation is infeasible, ZFitter provides a useful standalone approach to estimating acute viral infection times.

## ACKNOWLEDGEMENTS

This work was supported by funding to B.M. from the Swedish Research Council (2018-02381), and the US National Institute Of Allergy And Infectious Diseases, National Institutes of Health (UM1AI068618).

## AUTHOR CONTRIBUTIONS

Conceptualization, AP, MC, BM; Formal Analysis, AP, MC, BM; Investigation, AP; Visualization, AP; Writing – Original Draft, AP, MC, NS, DS, BM; Writing – Review & Editing, AP, MC, NS, DS, BM; Funding Acquisition, BM.

## Supplementary Note 1: Julia Implementation

~~~
using BioSequences
using Distributions: quantile, Beta
using StatsBase: countmap, median
ZF_cdf(p,a,b) = -log(quantile(Beta(a,b),1-p));
rate = 1.19e-5; #subs site ^-1 day ^-1
reader = open(FASTA.Reader, “ungapped-sequences.fasta”);
seqs = [sequence(record)) for record in reader];
close(reader);
counts = countmap(seqs);
N = length(seqs);
s = N - sum(values(counts). = = 1);
seqLen = median(length.(seqs));
#posterior interval incorporating (0.3, 1.0) prior
p = [0.025, 0.5, 0.975];
ZF_cdf.(p, s + 0.3, N - s + 1.0)./ (seqLen * rate)
~~~

## Supplementary Note 2: Python Implementation

~~~
from Bio import SeqIO
from statistics import median
from scipy.special import betaincinv
from math import log
def ZF_cdf(p, a, b):
return -log(betaincinv(a, b, 1 - p))
rate = 1.19e-5 #subs site ^-1 day ^-1
seqs = [str(r.seq) for r in SeqIO.parse(“ungapped-sequences.fasta”, “fasta”)]
counts = dict()
for s in seqs:
counts[s] = counts.get(s,0) +1
N = len(seqs)
s = N - sum([v = = 1 for v in counts.values()])
seqLen = median([len(s) for s in seqs])
#posterior interval incorporating (0.3, 1.0) prior
p = [0.025, 0.5, 0.975]
[ZF_cdf(p_i, s + 0.3, N - s + 1.0) / (seqLen * rate) for p_i in p]
~~~

## Supplementary Note 3: R Implementation

~~~
library(“Biostrings”)
library(“stats”)
ZF_cdf <- function(p, a, b) { -log(qbeta(1-p, a, b))}
rate = 1.19e-5 #subs site ^-1 day ^-1
seqs <- readDNAStringSet(“upgapped-sequences.fasta”, “fasta”)
t <- sort(table(seqs), decreasing = T)
N <- length(seqs)
s <- sum(t[t ! = 1])
seqLen <- median(Biostrings::width(seqs))
#posterior interval incorporating (0.3, 1.0) prior
p <- c(0.025, 0.5, 0.975)
ZF_cdf(p, s + 0.3, N - s + 1.0) / (seqLen * rate)
~~~

---

**Table S1.** ZFitter and Poisson-Fitter results for published Sanger SGA datasets from acute infection and associated metadata.

| Subject | Fiebig Stage | Subtype | Founders | Recombination | N | S | ZFitter median (95% CI) | Poisson-Fitter median (95% CI) | HD Poisson | Study |
| --- | --- | --- | --- | --- | --- | --- | --- | --- | --- | --- |
| 1006 | III | B | 1 | NA | 42 | 30 | 12 (6, 19) | 10 (5, 14) | True | Keele_2008 |
| 1054 | II | B | 1 | NA | 39 | 24 | 16 (10, 26) | 15 (9, 22) | True | Keele_2008 |
| 1056 | II | B | 1 | NA | 46 | 29 | 15 (9, 24) | 12 (7, 16) | True | Keele_2008 |
| 6240 | II | B | 1 | NA | 17 | 11 | 15 (6, 30) | 19 (11, 28) | True | Keele_2008 |
| 6244 | II | B | 1 | NA | 11 | 6 | 21 (8, 45) | 21 (7, 34) | True | Keele_2008 |
| 9010 | II | B | 1 | NA | 19 | 11 | 19 (9, 35) | 20 (11, 29) | True | Keele_2008 |
| 9014 | II | B | 1 | NA | 14 | 7 | 24 (11, 45) | 18 (9, 27) | True | Keele_2008 |
| 9015 | II | B | 1 | NA | 36 | 25 | 13 (7, 21) | 11 (7, 16) | True | Keele_2008 |
| 9017 | II | B | 1 | NA | 26 | 21 | 8 (3, 16) | 9 (3, 15) | True | Keele_2008 |
| 9019 | V | B | 1 | NA | 18 | 5 | 38 (21, 64) | 24 (17, 30) | True | Keele_2008 |
| 9020 | II | B | 1 | NA | 25 | 14 | 20 (11, 34) | 25 (18, 33) | True | Keele_2008 |
| 9021 | II | B | 1 | NA | 34 | 29 | 6 (2, 12) | 9 (3, 15) | True | Keele_2008 |
| 9023 | II | B | 1 | NA | 18 | 5 | 38 (21, 65) | 23 (12, 33) | True | Keele_2008 |
| 9024 | II | B | 1 | NA | 25 | 8 | 34 (20, 54) | 27 (18, 35) | True | Keele_2008 |
| 9025 | II | B | 1 | NA | 19 | 9 | 23 (12, 40) | 16 (8, 24) | True | Keele_2008 |
| 9028 | II | B | 1 | NA | 23 | 14 | 14 (7, 26) | 9 (4, 14) | True | Keele_2008 |
| 9032 | III | B | 1 | NA | 39 | 26 | 14 (8, 22) | 9 (6, 13) | True | Keele_2008 |
| 9033 | III | B | 1 | NA | 20 | 8 | 28 (15, 47) | 21 (12, 29) | True | Keele_2008 |
| 9075 | II | B | 1 | NA | 22 | 12 | 17 (9, 30) | 11 (5, 17) | True | Keele_2008 |
| 9077 | II | B | 1 | NA | 24 | 15 | 16 (8, 29) | 15 (7, 22) | True | Keele_2008 |
| 9079 | II | B | 1 | NA | 26 | 18 | 13 (6, 23) | 14 (7, 21) | True | Keele_2008 |
| 61792 | II | B | 1 | NA | 19 | 8 | 24 (13, 42) | 15 (8, 22) | True | Keele_2008 |
| 62130 | II | B | 1 | NA | 11 | 7 | 17 (6, 37) | 19 (6, 31) | True | Keele_2008 |
| 62357 | II | B | 1 | NA | 14 | 7 | 25 (11, 47) | 19 (11, 28) | True | Keele_2008 |
| 62995 | I | B | 1 | NA | 27 | 10 | 28 (17, 44) | 13 (7, 19) | True | Keele_2008 |
| 63054 | II | B | 1 | NA | 20 | 12 | 18 (8, 33) | 25 (16, 33) | True | Keele_2008 |
| 63396 | II | B | 1 | NA | 21 | 15 | 11 (4, 21) | 6 (2, 11) | True | Keele_2008 |
| PRB926 | II | B | 1 | NA | 14 | 7 | 24 (11, 47) | 27 (11, 43) | True | Keele_2008 |
| PRB931 | III | B | 1 | NA | 19 | 5 | 46 (25, 76) | 42 (31, 53) | True | Keele_2008 |
| PRB956 | II | B | 1 | NA | 27 | 16 | 18 (9, 30) | 15 (9, 21) | True | Keele_2008 |
| PRB958 | III | B | 1 | NA | 24 | 12 | 24 (13, 40) | 24 (17, 31) | True | Keele_2008 |
| PRB959 | II | B | 1 | NA | 32 | 21 | 15 (8, 25) | 13 (6, 19) | True | Keele_2008 |
| REJO | V | B | 1 | NA | 21 | 10 | 25 (13, 43) | 23 (13, 33) | True | Keele_2008 |
| SC05 | II | B | 1 | NA | 30 | 19 | 16 (8, 27) | 21 (15, 28) | True | Keele_2008 |
| SC11 | II | B | 1 | NA | 20 | 16 | 8 (3, 18) | 3 (0, 7) | True | Keele_2008 |
| SC20 | IV | B | 1 | NA | 43 | 27 | 16 (9, 25) | 14 (8, 19) | True | Keele_2008 |
| SC45 | II | B | 1 | NA | 29 | 15 | 22 (13, 36) | 18 (12, 24) | True | Keele_2008 |
| THRO | V | B | 1 | NA | 27 | 17 | 16 (8, 27) | 17 (9, 26) | True | Keele_2008 |
| TRJO | II | B | 1 | NA | 18 | 6 | 37 (20, 64) | 36 (22, 49) | True | Keele_2008 |
| TT28P | V | B | 1 | NA | 28 | 18 | 15 (8, 26) | 13 (8, 19) | True | Keele_2008 |
| TT29P | II | B | 1 | NA | 20 | 16 | 8 (3, 18) | 8 (2, 14) | True | Keele_2008 |
| TT34P | V | B | 1 | NA | 29 | 17 | 18 (10, 31) | 18 (11, 24) | True | Keele_2008 |
| WEAUd15 | II | B | 1 | NA | 44 | 22 | 23 (15, 34) | 23 (17, 29) | True | Keele_2008 |
| WITO | II | B | 1 | NA | 16 | 9 | 20 (9, 39) | 24 (12, 36) | True | Keele_2008 |
| Z02 | V | B | 1 | NA | 21 | 11 | 22 (11, 39) | 26 (17, 35) | True | Keele_2008 |
| Z05 | II | B | 1 | NA | 15 | 6 | 31 (15, 57) | 29 (19, 40) | True | Keele_2008 |
| Z13 | V | B | 1 | NA | 31 | 8 | 46 (29, 69) | 57 (45, 68) | True | Keele_2008 |
| Z20 | III | B | 1 | NA | 26 | 15 | 19 (10, 32) | 18 (12, 25) | True | Keele_2008 |
| Z23 | V | B | 1 | NA | 15 | 9 | 18 (8, 36) | 14 (7, 20) | True | Keele_2008 |
| Z27 | V | B | 1 | NA | 25 | 11 | 28 (16, 45) | 33 (22, 44) | True | Keele_2008 |
| Z32 | IV | B | 1 | NA | 10 | 6 | 19 (6, 41) | 20 (7, 33) | True | Keele_2008 |
| Z34 | III | B | 1 | NA | 18 | 11 | 18 (8, 33) | 28 (14, 42) | True | Keele_2008 |
| Z36 | VI | B | 1 | NA | 17 | 5 | 41 (22, 72) | 49 (36, 62) | True | Keele_2008 |
| 1018 | III | B | 1 | NA | 50 | 26 | 22 (14, 32) | 34 (24, 43) | False | Keele_2008 |
| 1053 | III | B | 1 | NA | 60 | 38 | 15 (10, 23) | 14 (6, 21) | False | Keele_2008 |
| 6248 | III | B | 1 | NA | 20 | 7 | 32 (17, 53) | 67 (25, 109) | False | Keele_2008 |
| 1001 | III | B | 1 | NA | 62 | 34 | 20 (13, 28) | 27 (20, 35) | False | Keele_2008 |
| RHPA | V | B | 1 | NA | 31 | 14 | 27 (16, 42) | 32 (21, 43) | True | Keele_2008 |
| 9022 | III | B | 1 | NA | 23 | 10 | 25 (14, 41) | 14 (9, 19) | True | Keele_2008 |
| 63215 | V | B | 1 | NA | 19 | 6 | 33 (18, 55) | 17 (9, 26) | True | Keele_2008 |
| TT35P | II | B | 1 | NA | 43 | 31 | 11 (6, 18) | 21 (7, 35) | False | Keele_2008 |
| 63358 | II | B | 1 | NA | 27 | 10 | 27 (16, 43) | 34 (24, 44) | True | Keele_2008 |
| 9031 | IV | B | 1 | NA | 20 | 4 | 44 (25, 72) | 37 (24, 50) | True | Keele_2008 |
| 9029 | II | B | 1 | NA | 22 | 7 | 32 (19, 53) | 20 (11, 29) | True | Keele_2008 |
| 1059 | III | B | 1 | NA | 39 | 17 | 28 (18, 41) | 41 (24, 58) | False | Keele_2008 |
| 12007 | II | B | 1 | NA | 25 | 8 | 34 (20, 54) | 27 (19, 36) | True | Keele_2008 |
| 1058 | IV | B | 1 | NA | 45 | 22 | 24 (15, 35) | 29 (20, 38) | False | Keele_2008 |
| SUMAd5 | II | B | 1 | NA | 35 | 23 | 14 (8, 24) | 13 (9, 17) | False | Keele_2008 |
| 1012 | III | B | 1 | NA | 43 | 23 | 21 (13, 31) | 31 (25, 38) | True | Keele_2008 |
| SC51 | V | B | 1 | NA | 33 | 21 | 16 (8, 26) | 23 (17, 28) | True | Keele_2008 |
| SC22 | II | B | 1 | NA | 38 | 21 | 19 (12, 30) | 30 (22, 38) | False | Keele_2008 |
| 9030 | II | B | 1 | NA | 19 | 7 | 28 (15, 48) | 27 (9, 45) | False | Keele_2008 |
| SC31 | IV | B | 1 | NA | 36 | 20 | 20 (12, 31) | 31 (18, 44) | False | Keele_2008 |
| CH40E | V | B | 1 | NA | 29 | 18 | 17 (9, 28) | 23 (17, 30) | True | Keele_2008 |
| CH77E | V | B | 1 | NA | 51 | 16 | 35 (25, 49) | 35 (29, 41) | True | Keele_2008 |
| CH58E | III | B | 1 | NA | 46 | 19 | 30 (20, 43) | 36 (28, 44) | False | Keele_2008 |
| MEMI | V | B | 1 | NA | 32 | 9 | 42 (27, 64) | 83 (68, 98) | False | Keele_2008 |
| Z33 | II | B | 1 | NA | 21 | 12 | 19 (10, 35) | 41 (31, 52) | True | Keele_2008 |
| 6247 | II | B | 2 | Yes | 32 | 26 | 8 (3, 15) | 25 (17, 33) | False | Keele_2008 |
| TT31P | II | B | 2 | Yes | 67 | 38 | 19 (13, 27) | 45 (39, 50) | False | Keele_2008 |
| Z31 | II | B | >1 | No | 17 | 2 | 72 (39, 128) | 111 (92, 130) | True | Keele_2008 |
| 63068 | II | B | 2 | Yes | 20 | 11 | 21 (10, 37) | NA | False | Keele_2008 |
| TT27P | IV | B | 3 | No | 38 | 24 | 15 (9, 25) | NA | False | Keele_2008 |
| 62615 | II | B | 3 | No | 28 | 13 | 22 (13, 36) | NA | False | Keele_2008 |
| Z35 | IV | B | 2 | Yes | 21 | 13 | 17 (8, 31) | NA | False | Keele_2008 |
| 9026 | III | B | 2 | No | 15 | 8 | 18 (8, 35) | NA | False | Keele_2008 |
| 9076 | III | B | 2 | No | 32 | 21 | 14 (8, 25) | NA | False | Keele_2008 |
| CH19E | V | B | >3 | Yes | 33 | 10 | 39 (25, 58) | NA | False | Keele_2008 |
| Z18 | V | B | 3 | Yes | 34 | 10 | 41 (26, 60) | NA | False | Keele_2008 |
| SC33 | II | B | 2 | No | 27 | 15 | 20 (11, 33) | NA | False | Keele_2008 |
| CAAN | V | B | >2 | Yes | 40 | 15 | 33 (21, 48) | NA | False | Keele_2008 |
| BORId9 | II | B | 5 | Yes | 29 | 10 | 35 (22, 55) | NA | False | Keele_2008 |
| 1051 | III | B | 4 | Yes | 50 | 29 | 18 (11, 27) | NA | False | Keele_2008 |
| Z30 | V | B | 2 | Yes | 30 | 3 | 76 (47, 119) | NA | False | Keele_2008 |
| Z16 | V | B | 5 | Yes | 19 | 2 | 75 (42, 131) | NA | False | Keele_2008 |
| Z03 | V | B | 3 | Yes | 22 | 0 | 188 (82, 516) | NA | False | Keele_2008 |
| PRB957 | II | B | 4 | No | 36 | 18 | 23 (14, 35) | NA | False | Keele_2008 |
| CH16E | V | B | 2 | Yes | 20 | 7 | 36 (20, 60) | NA | False | Keele_2008 |
| 12008 | II | B | 2 | No | 31 | 19 | 16 (9, 27) | NA | False | Keele_2008 |
| SC42 | IV | B | >3 | Yes | 25 | 0 | 192 (86, 520) | NA | False | Keele_2008 |
| Z10 | VI | B | >2 | Yes | 17 | 2 | 71 (39, 127) | NA | False | Keele_2008 |
| Z29 | V | B | >3 | Yes | 16 | 0 | 175 (72, 499) | NA | False | Keele_2008 |
| 4013171 | IV | B | >9 | Yes | 86 | 22 | 44 (33, 57) | 662 (525, 792) | False | Li_2010 |
| 4013211 | III | B | 2 | No | 30 | 13 | 28 (16, 43) | 54 (44, 64) | False | Li_2010 |
| 4013226 | II | B | 1 | NA | 33 | 18 | 20 (12, 33) | 15 (9, 21) | True | Li_2010 |
| 4013240 | II | B | 3 | Yes | 66 | 33 | 23 (16, 32) | 66 (56, 76) | False | Li_2010 |
| 4013242 | IV | B | 1 | NA | 37 | 16 | 28 (18, 42) | 16 (10, 22) | True | Li_2010 |
| 4013291 | V | B | 1 | NA | 25 | 0 | 192 (87, 522) | 61 (53, 70) | False | Li_2010 |
| 4013296 | II | B | 1 | NA | 25 | 8 | 37 (22, 59) | 34 (26, 42) | True | Li_2010 |
| 4013321 | II | B | 1 | NA | 49 | 24 | 24 (16, 35) | 39 (33, 45) | True | Li_2010 |
| 4013327 | IV | B | 1 | NA | 24 | 17 | 12 (6, 23) | 11 (5, 18) | True | Li_2010 |
| 4013383 | II | B | 2 | No | 70 | 49 | 12 (8, 18) | 554 (534, 572) | False | Li_2010 |
| 4013396 | IV | B | 1 | NA | 39 | 22 | 19 (12, 30) | 16 (11, 21) | True | Li_2010 |
| 4013419 | II | B | 3 | Yes | 78 | 40 | 22 (16, 30) | 548 (506, 591) | False | Li_2010 |
| 4013440 | II | B | 1 | NA | 30 | 10 | 37 (23, 56) | 23 (18, 28) | True | Li_2010 |
| 4013446 | III | B | 1 | NA | 23 | 14 | 17 (8, 30) | 24 (12, 35) | False | Li_2010 |
| 4013448 | II | B | 4 | Yes | 54 | 23 | 28 (19, 39) | 411 (367, 456) | False | Li_2010 |
| 4013448 | II | B | 4 | Yes | 54 | 23 | 28 (19, 39) | 411 (367, 456) | False | Li_2010 |
| AD75 | II | B | 1 | NA | 54 | 31 | 19 (12, 27) | 9 (6, 13) | True | Li_2010 |
| AD77 | V | B | 3 | NA | 40 | 7 | 58 (39, 84) | 84 (74, 95) | False | Li_2010 |
| AD83 | V | B | 3 | Yes | 44 | 18 | 30 (20, 43) | 66 (36, 95) | False | Li_2010 |
| HOBRO961 | II | B | 1 | NA | 42 | 24 | 19 (11, 29) | 17 (13, 22) | True | Li_2010 |
| INME0632 | II | B | 1 | NA | 46 | 29 | 16 (9, 24) | 12 (8, 16) | True | Li_2010 |
| 701010055 | II | B | 1 | NA | 28 | 16 | 19 (10, 32) | 15 (10, 21) | True | Li_2010 |
| 701010068 | IV | B | 7 | Yes | 89 | 5 | 52 (39, 70) | 688 (583, 792) | False | Li_2010 |
| 700010106 | II | B | 1 | NA | 40 | 15 | 32 (21, 47) | 13 (8, 18) | True | Li_2010 |
| 701010027 | V | B | 1 | NA | 27 | 9 | 37 (22, 58) | 42 (32, 52) | True | Li_2010 |
| 701010108 | V | B | 1 | NA | 35 | 16 | 26 (16, 40) | 39 (34, 45) | False | Li_2010 |
| 700010246 | IV | B | 1 | NA | 45 | 24 | 21 (13, 31) | 19 (13, 25) | True | Li_2010 |
| 700010238 | V | B | 3 | Yes | 38 | 12 | 38 (25, 55) | 161 (149, 174) | False | Li_2010 |

**Fig. ED1.**
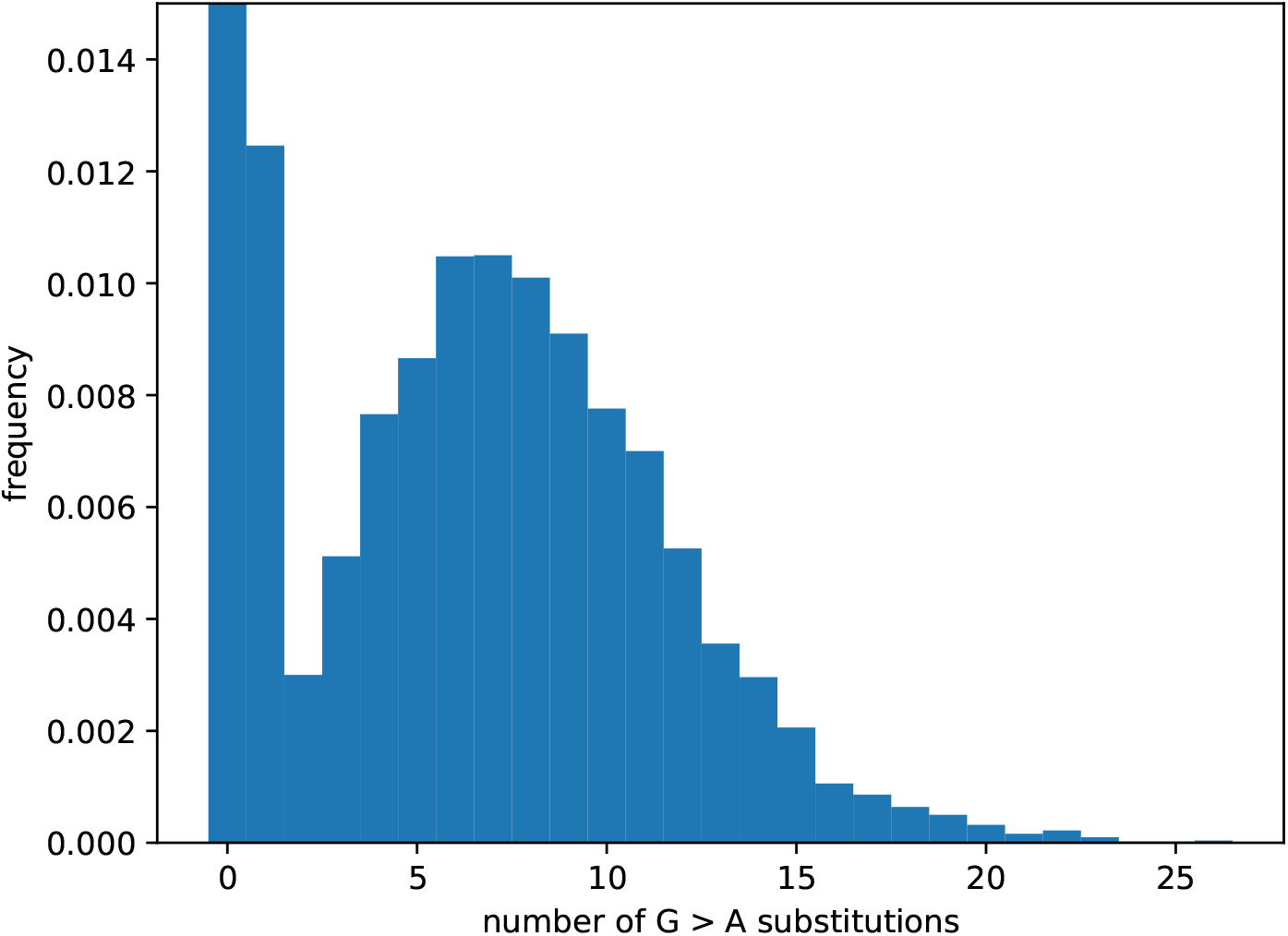
Histogram of G to A mutation counts for 50,000 simulated APOBEC events. The frequecy of the 0th bin (no G to A mutations) was 0.890.

## Bibliography

1. Peter B. Gilbert, Michal Juraska, Allan C. deCamp, Shelly Karuna, Srilatha Edupuganti, Nyaradzo Mgodi, Deborah J. Donnell, Carter Bentley, Nirupama Sista, Philip Andrew, Abby Isaacs, Yunda Huang, Lily Zhang, Edmund Capparelli, Nidhi Kochar, Jing Wang, Susan H. Eshleman, Kenneth H. Mayer, Craig A. Magaret, John Hural, James G. Kublin, Glenda Gray, David C. Montefiori, Margarita M. Gomez, David N. Burns, Julie McElrath, Julie Ledgerwood, Barney S. Graham, John R. Mascola, Myron Cohen, and Lawrence Corey. Basis and Statistical Design of the Passive HIV-1 Antibody Mediated Prevention (AMP) Test-of-Concept Efficacy Trials. Statistical Communications in Infectious Diseases, 9(1), January 2017. ISSN 1948-4690. doi: 10.1515/scid-2016-0001.

2. B. F. Keele, E. E. Giorgi, J. F. Salazar-Gonzalez, J. M. Decker, K. T. Pham, M. G. Salazar, C. Sun, T. Grayson, S. Wang, H. Li, X. Wei, C. Jiang, J. L. Kirchherr, F. Gao, J. A. Anderson, L.-H. Ping, R. Swanstrom, G. D. Tomaras, W. A. Blattner, P. A. Goepfert, J. M. Kilby, M. S. Saag, E. L. Delwart, M. P. Busch, M. S. Cohen, D. C. Montefiori, B. F. Haynes, B. Gaschen, G. S. Athreya, H. Y. Lee, N. Wood, C. Seoighe, A. S. Perelson, T. Bhattacharya, B. T. Korber, B. H. Hahn, and G. M. Shaw. Identification and characterization of transmitted and early founder virus envelopes in primary HIV-1 infection. Proceedings of the National Academy of Sciences, 105(21):7552–7557, May 2008. ISSN 0027-8424, 1091-6490. doi: 10.1073/pnas.0802203105.

3. M.-R. Abrahams, J. A. Anderson, E. E. Giorgi, C. Seoighe, K. Mlisana, L.-H. Ping, G. S. Athreya, F. K. Treurnicht, B. F. Keele, N. Wood, J. F. Salazar-Gonzalez, T. Bhattacharya, H. Chu, I. Hoffman, S. Galvin, C. Mapanje, P. Kazembe, R. Thebus, S. Fiscus, W. Hide, M. S. Cohen, S. Abdool Karim, B. F. Haynes, G. M. Shaw, B. H. Hahn, B. T. Korber, R. Swanstrom, C. Williamson, CAPRISA Acute Infection Study Team, and Center for HIV-AIDS Vaccine Immunology Consortium. Quantitating the multiplicity of infection with human immunodeficiency virus type 1 subtype C reveals a non-poisson distribution of transmitted variants. Journal of Virology, 83(8):3556–3567, April 2009. ISSN 1098-5514. doi: 10.1128/JVI.02132-08.

4. Richard E. Haaland, Paulina A. Hawkins, Jesus Salazar-Gonzalez, Amber Johnson, Amanda Tichacek, Etienne Karita, Olivier Manigart, Joseph Mulenga, Brandon F. Keele, George M. Shaw, Beatrice H. Hahn, Susan A. Allen, Cynthia A. Derdeyn, and Eric Hunter. Inflammatory Genital Infections Mitigate a Severe Genetic Bottleneck in Heterosexual Transmission of Subtype A and C HIV-1. PLOS Pathogens, 5(1):e1000274, January 2009. ISSN 1553-7374. doi: 10.1371/journal.ppat.1000274.

5. Katharine J. Bar, Hui Li, Annie Chamberland, Cecile Tremblay, Jean Pierre Routy, Truman Grayson, Chuanxi Sun, Shuyi Wang, Gerald H. Learn, Charity J. Morgan, Joseph E. Schumacher, Barton F. Haynes, Brandon F. Keele, Beatrice H. Hahn, and George M. Shaw. Wide variation in the multiplicity of HIV-1 infection among injection drug users. Journal of Virology, 84(12):6241–6247, June 2010. ISSN 1098-5514. doi: 10.1128/JVI.00077-10.

6. Hui Li, Katharine J. Bar, Shuyi Wang, Julie M. Decker, Yalu Chen, Chuanxi Sun, Jesus F. Salazar-Gonzalez, Maria G. Salazar, Gerald H. Learn, Charity J. Morgan, Joseph E. Schumacher, Peter Hraber, Elena E. Giorgi, Tanmoy Bhattacharya, Bette T. Korber, Alan S. Perelson, Joseph J. Eron, Myron S. Cohen, Charles B. Hicks, Barton F. Haynes, Martin Markowitz, Brandon F. Keele, Beatrice H. Hahn, and George M. Shaw. High Multiplicity Infection by HIV-1 in Men Who Have Sex with Men. PLoS pathogens, 6(5):e1000890, May 2010. ISSN 1553-7374. doi: 10.1371/journal.ppat.1000890.

7. Ruy M. Ribeiro, Li Qin, Leslie L. Chavez, Dongfeng Li, Steven G. Self, and Alan S. Perelson. Estimation of the Initial Viral Growth Rate and Basic Reproductive Number during Acute HIV-1 Infection. Journal of Virology, 84(12):6096–6102, June 2010. ISSN 0022-538X. doi: 10.1128/JVI.00127-10.

8. M. Slatkin and R. R. Hudson. Pairwise comparisons of mitochondrial DNA sequences in stable and exponentially growing populations. Genetics, 129(2):555–562, October 1991. ISSN 0016-6731, 1943-2631.

9. Raj Shankarappa, Joseph B. Margolick, Stephen J. Gange, Allen G. Rodrigo, David Upchurch, Homayoon Farzadegan, Phalguni Gupta, Charles R. Rinaldo, Gerald H. Learn, Xi He, Xiao-Li Huang, and James I. Mullins. Consistent Viral Evolutionary Changes Associated with the Progression of Human Immunodeficiency Virus Type 1 Infection. Journal of Virology, 73(12):10489–10502, December 1999. ISSN 0022-538X.

10. Art F. Y. Poon, Rachel A. McGovern, Theresa Mo, David J. H. F. Knapp, Bluma Brenner, Jean-Pierre Routy, Mark A. Wainberg, and P. Richard Harrigan. Dates of HIV infection can be estimated for seroprevalent patients by coalescent analysis of serial next-generation sequencing data. AIDS, 25(16):2019–2026, October 2011. ISSN 0269-9370. doi: 10.1097/QAD.0b013e32834b643c.

11. Cynthia Gay, Oliver Dibben, Jeffrey A. Anderson, Andrea Stacey, Ashley J. Mayo, Philip J. Norris, JoAnn D. Kuruc, Jesus F. Salazar-Gonzalez, Hui Li, Brandon F. Keele, Charles Hicks, David Margolis, Guido Ferrari, Barton Haynes, Ronald Swanstrom, George M. Shaw, Beatrice H. Hahn, Joseph J. Eron, Persephone Borrow, and Myron S. Cohen. Cross-Sectional Detection of Acute HIV Infection: Timing of Transmission, Inflammation and Antiretroviral Therapy. PLOS ONE, 6(5):e19617, May 2011. ISSN 1932-6203. doi: 10.1371/journal.pone.0019617.

12. Manon Ragonnet-Cronin, Stéphane Aris-Brosou, Isabelle Joanisse, Harriet Merks, Dominic Vallée, Kyna Caminiti, Michael Rekart, Mel Krajden, Darrel Cook, John Kim, Laurie Malloch, Paul Sandstrom, and James Brooks. Genetic Diversity as a Marker for Timing Infection in HIV-Infected Patients: Evaluation of a 6-Month Window and Comparison With BED. TheJournal of Infectious Diseases, 206(5):756–764, September 2012. ISSN 0022-1899. doi: 10.1093/infdis/jis411.

13. Massimo Ciccozzi, Alessandra Lo Presti, Mauro Andreotti, Sandro Mancinelli, Susanna Ceffa, Clementina Maria Galluzzo, Ersilia Buonomo, Richard Luhanga, Haswell Jere, Eleonora Cella, Paola Scarcella, Marco Mirra, Maria Cristina Marazzi, Stefano Vella, Leonardo Palombi, and Marina Giuliano. Viral Sequence Analysis of HIV-Positive Women and Their Infected Children: Insight on the Timing of Infection and on the Transmission Network. AIDS Research and Human Retroviruses, 30(10):1010–1015, August 2014. ISSN 0889-2229. doi: 10.1089/aid.2014.0143.

14. Vadim Puller, Richard Neher, and Jan Albert. Estimating time of HIV-1 infection from next-generation sequence diversity. PLOS Computational Biology, 13(10):e1005775, October 2017. ISSN 1553-7358. doi: 10.1371/journal.pcbi.1005775.

15. Elena E. Giorgi, Bob Funkhouser, Gayathri Athreya, Alan S. Perelson, Bette T. Korber, and Tanmoy Bhattacharya. Estimating time since infection in early homogeneous HIV-1 samples using a poisson model. BMC bioinformatics, 11:532, October 2010. ISSN 1471-2105. doi: 10.1186/1471-2105-11-532.

16. Ha Youn Lee, Elena E. Giorgi, Brandon F. Keele, Brian Gaschen, Gayathri S. Athreya, Jesus F. Salazar-Gonzalez, Kimmy T. Pham, Paul A. Goepfert, J. Michael Kilby, Michael S. Saag, Eric L. Delwart, Michael P. Busch, Beatrice H. Hahn, George M. Shaw, Bette T. Korber, Tanmoy Bhattacharya, and Alan S. Perelson. Modeling sequence evolution in acute HIV-1 infection. Journal of Theoretical Biology, 261(2):341–360, November 2009. ISSN 0022-5193. doi: 10.1016/j.jtbi.2009.07.038.

17. Ann M. Sheehy, Nathan C. Gaddis, and Michael H. Malim. The antiretroviral enzyme APOBEC3G is degraded by the proteasome in response to HIV-1 Vif. Nature Medicine, 9(11):1404–1407, November 2003. ISSN 1546-170X. doi: 10.1038/nm945.

18. Heather L Wiegand, Brian P Doehle, Hal P Bogerd, and Bryan R Cullen. A second human antiretroviral factor, APOBEC3F, is suppressed by the HIV-1 and HIV-2 Vif proteins. The EMBO Journal, 23(12):2451–2458, June 2004. ISSN 0261-4189. doi: 10.1038/sj.emboj.7600246.

19. Rupert C. L. Beale, Svend K. Petersen-Mahrt, Ian N. Watt, Reuben S. Harris, Cristina Rada, and Michael S. Neuberger. Comparison of the Differential Context-dependence of DNA Deamination by APOBEC Enzymes: Correlation with Mutation Spectra in Vivo. Journal of Molecular Biology, 337(3):585–596, March 2004. ISSN 0022-2836. doi: 10.1016/j.jmb.2004.01.046.

20. Viviana Simon, Veronique Zennou, Deya Murray, Yaoxing Huang, David D. Ho, and Paul D. Bieniasz. Natural Variation in Vif: Differential Impact on APOBEC3G/3F and a Potential Role in HIV-1 Diversification. PLOS Pathogens, 1(1):e6, July 2005. ISSN 1553-7374. doi: 10.1371/journal.ppat.0010006.

21. Andrew E. Armitage, Koen Deforche, Chih-hao Chang, Edmund Wee, Beatrice Kramer, John J. Welch, Jan Gerstoft, Lars Fugger, Andrew McMichael, Andrew Rambaut, and Astrid K. N. Iversen. APOBEC3G-Induced Hypermutation of Human Immunodeficiency Virus Type-1 Is Typically a Discrete “All or Nothing” Phenomenon. PLOS Genetics, 8(3): e1002550, March 2012. ISSN 1553-7404. doi: 10.1371/journal.pgen.1002550.

22. Rebecca A. Russell, Michael D. Moore, Wei-Shau Hu, and Vinay K. Pathak. APOBEC3G induces a hypermutation gradient: purifying selection at multiple steps during HIV-1 replication results in levels of G-to-A mutations that are high in DNA, intermediate in cellular viral RNA, and low in virion RNA. Retrovirology, 6:16, February 2009. ISSN 1742-4690. doi: 10.1186/1742-4690-6-16.

23. Holly A. Sadler, Mark D. Stenglein, Reuben S. Harris, and Louis M. Mansky. APOBEC3G Contributes to HIV-1 Variation through Sublethal Mutagenesis. Journal of Virology, 84(14): 7396–7404, July 2010. ISSN 0022-538X, 1098-5514. doi: 10.1128/JVI.00056-10.

24. Elena E. Giorgi, Hui Li, Tanmoy Bhattacharya, George M. Shaw, and Bette Korber. Estimating the Timing of Early Simian-Human Immunodeficiency Virus Infections: a Comparison between Poisson Fitter and BEAST. mBio, 11(2), March 2020. ISSN 2150-7511. doi: 10.1128/mBio.00324-20.

25. Ch Julián Villabona-Arenas, Matthew Hall, Katrina A. Lythgoe, Stephen G. Gaffney, Roland R. Regoes, Stéphane Hué, and Katherine E. Atkins. Number of HIV-1 founder variants is determined by the recency of the source partner infection. Science, 369(6499): 103–108, July 2020. ISSN 0036-8075, 1095-9203. doi: 10.1126/science.aba5443.

26. Tanzy M. T. Love, Sung Yong Park, Elena E. Giorgi, Wendy J. Mack, Alan S. Perelson, and Ha Youn Lee. SPMM: estimating infection duration of multivariant HIV-1 infections. Bioinformatics, 32(9):1308–1315, May 2016. ISSN 1367-4803. doi: 10.1093/bioinformatics/btv749.

27. Eric Lewitus and Morgane Rolland. A non-parametric analytic framework for within-host viral phylogenies and a test for HIV-1 founder multiplicity. Virus Evolution, 5(2), July 2019. doi: 10.1093/ve/vez044.

28. Raabya Rossenkhan, Morgane Rolland, Jan P. L. Labuschagne, Roux-Cil Ferreira, Craig A. Magaret, Lindsay N. Carpp, Frederick A. Matsen Iv, Yunda Huang, Erika E. Rudnicki, Yuanyuan Zhang, Nonkululeko Ndabambi, Murray Logan, Ted Holzman, Melissa-Rose Abrahams, Colin Anthony, Sodsai Tovanabutra, Christopher Warth, Gordon Botha, David Matten, Sorachai Nitayaphan, Hannah Kibuuka, Fred K. Sawe, Denis Chopera, Leigh Anne Eller, Simon Travers, Merlin L. Robb, Carolyn Williamson, Peter B. Gilbert, and Paul T. Edlefsen. Combining Viral Genetics and Statistical Modeling to Improve HIV-1 Time-of-infection Estimation towards Enhanced Vaccine Efficacy Assessment. Viruses, 11(7), 2019. ISSN 1999-4915. doi: 10.3390/v11070607.

29. Andrew Gelman and Eric Loken. The Statistical Crisis in Science. American Scientist, 102: 460, November 2014. doi: 10.1511/2014.111.460.

30. M. Hasegawa, H. Kishino, and T. Yano. Dating of the human-ape splitting by a molecular clock of mitochondrial DNA. Journal of Molecular Evolution, 22(2):160–174, 1985. ISSN 0022-2844. doi: 10.1007/BF02101694.

31. Elise Landais, Ben Murrell, Bryan Briney, Sasha Murrell, Kimmo Rantalainen, Zachary T. Berndsen, Alejandra Ramos, Lalinda Wickramasinghe, Melissa Laird Smith, Kemal Eren, Natalia de Val, Mengyu Wu, Audrey Cappelletti, Jeffrey Umotoy, Yolanda Lie, Terri Wrin, Paul Algate, Po-Ying Chan-Hui, Etienne Karita, Andrew B. Ward, Ian A. Wilson, Dennis R. Burton, Davey Smith, Sergei L. Kosakovsky Pond, and Pascal Poignard. HIV Envelope Glycoform Heterogeneity and Localized Diversity Govern the Initiation and Maturation of a V2 Apex Broadly Neutralizing Antibody Lineage. Immunity, 47(5):990–1003.e9, November 2017. ISSN 1074-7613. doi: 10.1016/j.immuni.2017.11.002.

32. Eberhard W. Fiebig, David J. Wright, Bhupat D. Rawal, Patricia E. Garrett, Richard T. Schumacher, Lorraine Peddada, Charles Heldebrant, Richard Smith, Andrew Conrad, Steven H. Kleinman, and Michael P. Busch. Dynamics of HIV viremia and antibody seroconversion in plasma donors: implications for diagnosis and staging of primary HIV infection. AIDS (London, England), 17(13):1871–1879, September 2003. ISSN 0269-9370. doi: 10.1097/00002030-200309050-00005.

33. Alexei J. Drummond and Andrew Rambaut. BEAST: Bayesian evolutionary analysis by sampling trees. BMC Evolutionary Biology, 7(1):214, November 2007. ISSN 1471-2148. doi: 10.1186/1471-2148-7-214.

34. R. A. Koup, J. T. Safrit, Y. Cao, C. A. Andrews, G. McLeod, W. Borkowsky, C. Farthing, and D. D. Ho. Temporal association of cellular immune responses with the initial control of viremia in primary human immunodeficiency virus type 1 syndrome. Journal of Virology, 68 (7):4650–4655, July 1994. ISSN 0022-538X. doi: 10.1128/JVI.68.7.4650-4655.1994.

35. P. Borrow, H. Lewicki, B. H. Hahn, G. M. Shaw, and M. B. Oldstone. Virus-specific CD8 + cytotoxic T-lymphocyte activity associated with control of viremia in primary human immunodeficiency virus type 1 infection. Journal of Virology, 68(9):6103–6110, September 1994. ISSN 0022-538X, 1098-5514.

36. Yi Liu, John McNevin, Jianhong Cao, Hong Zhao, Indira Genowati, Kim Wong, Sherry McLaughlin, Matthew D. McSweyn, Kurt Diem, Claire E. Stevens, Janine Maenza, Hongxia He, David C. Nickle, Daniel Shriner, Sarah E. Holte, Ann C. Collier, Lawrence Corey, M. Juliana McElrath, and James I. Mullins. Selection on the Human Immunodeficiency Virus Type 1 Proteome following Primary Infection. Journal of Virology, 80(19):9519–9529, October 2006. ISSN 0022-538X, 1098-5514. doi: 10.1128/JVI.00575-06.

37. Nilu Goonetilleke, Michael K.P. Liu, Jesus F. Salazar-Gonzalez, Guido Ferrari, Elena Giorgi, Vitaly V. Ganusov, Brandon F. Keele, Gerald H. Learn, Emma L. Turnbull, Maria G. Salazar, Kent J. Weinhold, Stephen Moore, Norman Letvin, Barton F. Haynes, Myron S. Cohen, Peter Hraber, Tanmoy Bhattacharya, Persephone Borrow, Alan S. Perelson, Beatrice H. Hahn, George M. Shaw, Bette T. Korber, and Andrew J. McMichael. The first T cell response to transmitted/founder virus contributes to the control of acute viremia in HIV-1 infection. The Journal of Experimental Medicine, 206(6):1253–1272, June 2009. ISSN 0022-1007. doi: 10.1084/jem.20090365.

38. Hongshuo Song, Elena E. Giorgi, Vitaly V. Ganusov, Fangping Cai, Gayathri Athreya, Hyejin Yoon, Oana Carja, Bhavna Hora, Peter Hraber, Ethan Romero-Severson, Chunlai Jiang, Xiaojun Li, Shuyi Wang, Hui Li, Jesus F. Salazar-Gonzalez, Maria G. Salazar, Nilu Goonetilleke, Brandon F. Keele, David C. Montefiori, Myron S. Cohen, George M. Shaw, Beatrice H. Hahn, Andrew J. McMichael, Barton F. Haynes, Bette Korber, Tanmoy Bhattacharya, and Feng Gao. Tracking HIV-1 recombination to resolve its contribution to HIV-1 evolution in natural infection. Nature Communications, 9(1):1928, May 2018. ISSN 2041-1723. doi: 10.1038/s41467-018-04217-5.

39. Darren P. Martin, Ben Murrell, Michael Golden, Arjun Khoosal, and Brejnev Muhire. RDP4: Detection and analysis of recombination patterns in virus genomes. Virus Evolution, 1(1), March 2015. doi: 10.1093/ve/vev003.

